# PRDX6 dictates ferroptosis sensitivity by directing cellular selenium mobilization

**DOI:** 10.1101/2024.06.04.595693

**Authors:** Junya Ito, Toshitaka Nakamura, Takashi Toyama, Sebastian Doll, Mirai Suzuki, Weijia Zhang, Jiashuo Zheng, Dietrich Trümbach, Naoya Yamada, Koya Ono, Adam Wahida, Yoshiro Saito, Kiyotaka Nakagawa, Eikan Mishima, Marcus Conrad

## Abstract

Selenium-dependent glutathione peroxidase 4 (GPX4) is the guardian of ferroptosis and prevents unrestrained (phospho)lipid peroxidation by directly reducing phospholipid hydroperoxides (PLOOH) to their corresponding alcohols. However, it remains unclear whether other phospholipid peroxidases can also contribute to ferroptosis prevention, albeit to a varying degree. Here we show that cells lacking GPX4 still exhibit substantial PLOOH reduction capacity, arguing for the presence of alternative PLOOH peroxidases. By scrutinizing potential candidates, we showed that while overexpression of peroxiredoxin 6 (PRDX6), a thiol-specific antioxidant enzyme with reported PLOOH-reducing activity, failed to prevent ferroptosis, its genetic loss markedly sensitizes cancer cells to ferroptosis. Mechanistically, we uncover that PRDX6 facilitates intracellular selenium handling, which is crucial for selenium incorporation into selenoproteins, including GPX4. Consequently, PRDX6 modulates GPX4 expression, thereby dictating the sensitivity of cells to undergo ferroptosis. Our study highlights PRDX6 as a critical factor in ferroptosis prevention by directing cellular selenium mobilization.

## Introduction

Ferroptosis, a regulated cell death modality hallmarked by unrestrained iron-dependent (phospho)lipid peroxidation^1^, has emerged as an important facet of redox cell biology and a promising target for treating human disease^2–4^. Notably, its potential efficacy against (chemo)therapy-resistant and metastasizing cancers positions ferroptosis as a highly promising target for future therapeutic approaches^5–7^. Consequently, the search for viable therapeutic targets to induce ferroptosis has taken center stage in efforts to combat these therapy-resistant cancers^7^.

Cells have developed various defense systems capable of detoxifying deleterious phospholipid hydroperoxides (PLOOH) to avert ferroptosis^2,8,9^. Among these, selenium-dependent glutathione peroxidase 4 (GPX4) is considered crucial to prevent ferroptosis by directly reducing PLOOH to their corresponding alcohols at the expense of glutathione (GSH)^10,11^. Since GPX4 belongs to the selenoprotein family, which is characterized by proteins carrying at least one selenocysteine (Sec) usually in their active site, this residue is critical for enabling full GPX4 enzymatic activity and protection from peroxide-inducible, irreversible overoxidation and enzyme inactivation^12^. In addition to the GSH/GPX4 axis, a series of genetic screens have unveiled alternative ferroptosis surveillance systems, such as the NAD(P)H/ferroptosis suppressor protein-1 (FSP1)/ubiquinone or vitamin K system^13–15^, di-/tetrahydrobiopterin/GTP cyclohydrolase 1 (GCH1)/dihydrofolate reductase (DHFR) system^16,17^ and 7-dihydrocholesterol reductase (DHCR7)^18–20^, that can act as backup systems for GPX4 at least in certain cellular contexts. Unlike GPX4, these systems protect against the lipid peroxidation chain reaction by reducing phospholipid peroxyl radicals to phospholipid hydroperoxides that require additional steps for their decomposition.

Akin to GPX4, selenium-independent peroxiredoxin 6 (PRDX6), one of six thiol-specific antioxidant protein family members, has been reported to directly reduce PLOOH to their corresponding alcohols^9,21,22^. Interestingly, PRDX6 acts as a bifunctional enzyme with glutathione peroxidase and phospholipase A2 (PLA_2_) activity^23^. Its peroxidase activity depends on the catalytic Cys at position 47 and uses the cofactor GSH as the physiological reductant^24^. Earlier work suggested that genetic perturbation of PRDX6 renders cancer cells more vulnerable to ferroptosis^25–27^. In addition, PRDX6 has been repeatedly identified in genetic screens or gene cluster analyses aimed at identifying genes impinging on ferroptosis regulation^17,28^. However, the mechanisms by which PRDX6 modulate ferroptosis susceptibility or its impact on PLOOH reduction remain poorly understood.

In this study, we set out to unravel the purported role of PRDX6 in ferroptosis regulation by preventing uncontrolled (phospho)lipid peroxidation. Contrary to our expectations, the role of PRDX6 in determining ferroptosis sensitivity did not relate to intracellular PLOOH reduction or the PLA_2_ activity but rather to a crucial involvement in the biosynthesis of selenoproteins, including GPX4. These findings underscore PRDX6 as a pivotal player participating in intracellular selenium handling and mobilization, thereby closing a gap in our understanding of how PRDX6 impacts cellular susceptibility to ferroptosis.

## Results

### PCOOH reduction rates are unaffected by GPX4 deficiency in cells and liver tissue

To assess the exact contribution of GPX4 in PLOOH reduction, we established a mass spectrometry-based method to evaluate the rate of PLOOH reduction using cell lysate samples (**Fig. 1a**). Hereby, synthesized deuterium-labelled phosphatidylcholine hydroperoxide (PCOOH-*d*_9_; prepared as shown in **Fig. S1**) is incubated with cell lysates for 1 h, after which the rate of PCOOH reduction is evaluated using liquid chromatography-tandem mass spectrometry (LC-MS/MS). Incubation in the absence of cell lysates (i.e., blank group) showed minimal PCOOH reduction, whereas cell lysates collected from human fibrosarcoma HT-1080 cells efficiently reduced PCOOH (∼ 70% reduction) (**Fig. 1b**). On the contrary, cell lysates from *GPX4* knockout (KO) HT-1080 cells showed a significant decrease in PCOOH-reducing ability by ∼ 10% compared to WT lysates, which could be rescued by overexpression of GPX4. Interestingly, the ability to reduce PCOOH was preserved in *GPX4* KO cells (∼ 65% PCOOH reduction). We confirmed this residual PCOOH reducing ability also in Pfa1 cells, a widely used mouse embryonic fibroblasts in ferroptosis research, where treatment of cells with 4-hydroxytamoxifen (TAM) induces *Gpx4* deletion and ferroptosis^29^ (**Fig. S2**). In line with these *in vitro* findings, PCOOH levels were not different between liver tissue samples from control and hepatocyte-specific *Gpx4* KO mice maintained on a low vitamin E diet, which causes overt ferroptotic cell death in hepatocytes^13^ (**Fig. 1c**), suggesting that GPX4 may not be the prime enzyme responsible for overall cellular PCOOH reduction. Moreover, the presence of an alternative and GPX4-independent PCOOH reducing pathway is implied. Consequently, we shifted our focus to PRDX6, a putative alternative enzyme harboring PLOOH-reducing activity^21,22^.

**Figure 1.**
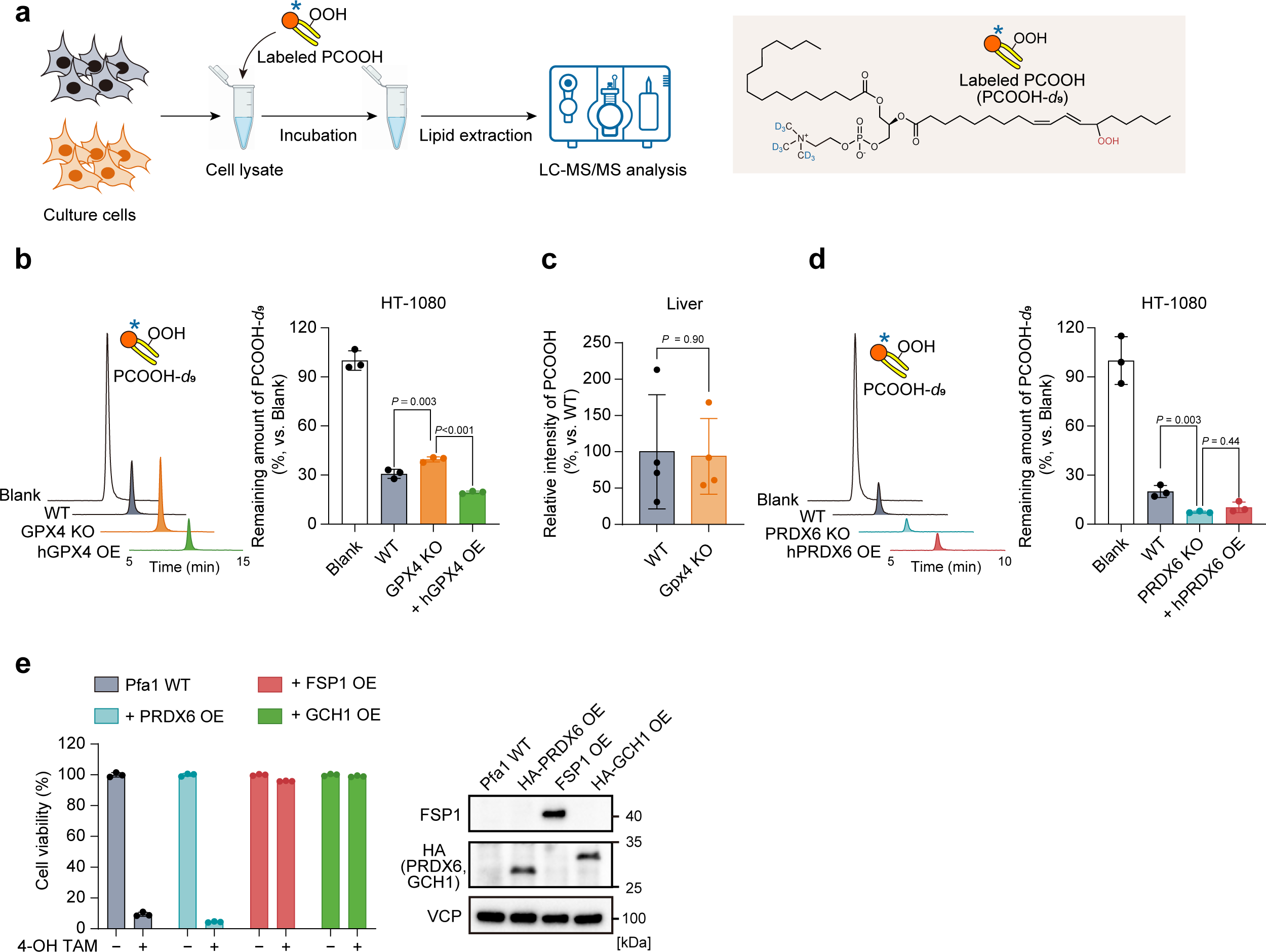
| PCOOH reducing capacity in cells lacking either GPX4 or PRDX6 **(a)** Scheme for the assessment of PCOOH reducing capacity using whole cell lysate samples. After the incubation of deuterium-labelled phosphatidylcholine hydroperoxide (PCOOH-*d*_9_) with cell lysate, the remaining amount of PCOOH-*d*_9_ was analyzed by LC-MS/MS. **(b)** PCOOH reducing capacity using cell lysates collected from WT, *GPX4* knockout (KO) and *GPX4* KO HT1080 cells overexpressing (OE) hGPX4. The chromatogram (left) and relative value of the remaining PCOOH-*d*_9_ (right) are shown. **(c)** Tissue PCOOH levels in the liver of control and hepatocyte-specific *Gpx4* KO mice. The mean value in the control group was taken as 100%. **(d)** PCOOH reducing capacity using cell lysate collected from WT, *PRDX6* KO and *PRDX6* KO HT1080 cells overexpressing of hPRDX6. **(e)** Cell viability of Pfa1 cells stably overexpressing HA-mPRDX6, hFSP1 or HA-hGCH1. Viability was measured 72 h after 4-hydroxytamoxifen (TAM) treatment to deplete *Gpx4*. Valosin-containing protein (VCP) was used as a loading control. Data is mean ± s.d. of n = 3 (b, d, e) and n = 4 (c). ANOVA (Tukey) in (b and d). t-test in (c). The illustration in (a) was generated using BioRender.com.

### Overexpression of PRDX6 fails to protect against ferroptosis

We next explored the PCOOH reductive capacity of PRDX6 in cells using our established PCOOH-*d*_9_ and LS-MS/MS method. This experiment showed that cell lysates collected from *PRDX6* KO HT-1080 cells also had a preserved PCOOH-reductive capacity (**Fig. 1d**). Moreover, cell lysates collected from *PRDX6 KO* cells and hPRDX6-reconstituted cells showed comparable PCOOH reduction capacity, suggesting that PRDX6 also does not mainly contribute to the overall cellular PCOOH reduction. Furthermore, in contrast to other ferroptosis suppressors, such as FSP1 and GCH1^14,16^, overexpression of PRDX6 consistently did not protect cells against ferroptosis induced by TAM-induced *Gpx4* deletion in Pfa1 cells (**Fig. 1e**). Hence, we concluded that overexpression of PRDX6, unlike GPX4, does not significantly increase the cellular PLOOH reducing capacity nor fails to prevent ferroptosis.

### PRDX6 deletion decreases GPX4 expression and sensitizes cancer cells to ferroptosis

To further investigate the potential role of PRDX6 in ferroptosis regulation, we examined its impact on GPX4 expression. Remarkably, GPX4 protein expression was considerably reduced in *PRDX6* KO HT-1080 cells (**Fig. 2a**), in contrast to the deletion of any of the other PRDX family members (PRDX1-5)^30^ (**Fig. 2b**). Similar observations were made across various cancer cell lines (**Fig. 2c**). Our results were supported by a significant correlation between PRDX6 and GPX4 protein expression across a comprehensive proteomic dataset (www.depmap.org) containing 375 cancer cell lines (**Fig. 2d**). To ascertain a functional consequence, we next evaluated ferroptosis susceptibility in cells lacking *PRDX6. PRDX6* KO HT1080 cells exhibited an increased ferroptosis sensitivity compared to WT cells when treated with a wide variety of ferroptosis inducers and sensitizers^31^, such as the GPX4 inhibitor (*1S,3R*)-RSL (RSL3)^11,32^; the cystine/glutamate antiporter (xCT, SLC7A11) inhibitor erastin^1^; the GSH biosynthesis inhibitor BSO and several FSP1 inhibitors (iFSP1^14^, viFSP1^33^ and icFSP1^34^) (**Fig. 2e**). The increased ferroptosis sensitivity in *PRDX6* KO HT-1080 cells was entirely restored by the ferroptosis inhibitor liproxstatin-1 (Lip-1)^10^. In addition, *PRDX6* KO consistently increased sensitivity to ferroptosis induced by RSL3 and erastin in various cancer cell lines (**Fig. 2f**), underscoring the important role of PRDX6 in modulating GPX4 expression and ensuing cells’ ferroptosis susceptibility.

**Figure 2.**
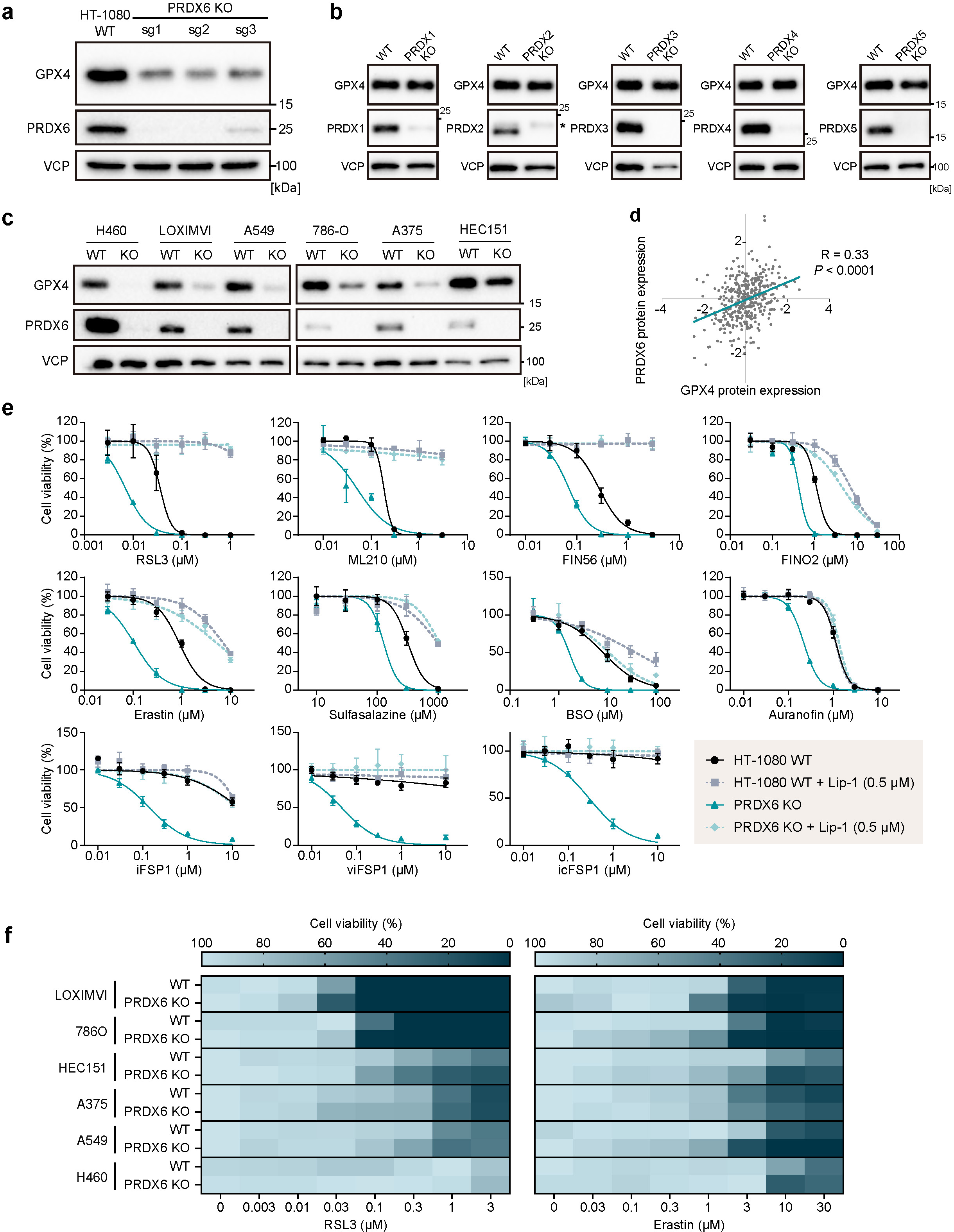
| Deletion of *PRDX6* decreases GPX4 protein expression and sensitizes cells to ferroptosis **(a)** Immunoblot analysis of GPX4 and PRDX6 in wild-type (WT) and *PRDX6* knockout (KO) HT-1080 cells. **(b)** Immunoblot analysis of GPX4 and PRDX1-5 in wild-type (WT) and *PRDX1-5* KO HT-1080 cells. For the PRDX2 immunoblot, the asterisk indicates a non-specific band. **(c)** Immunoblot analysis of WT and *PRDX6* KO NCI-H460, LOXIMVI, A549, 786-O, A375 and HEC151 cells. **(d)** Correlation of GPX4 and PRDX6 expression in proteomics data set (depmap; version v23Q4). **(e)** Viability of WT and *PRDX6* KO HT-1080 cells treated with various ferroptosis inducers and sensitizers. RSL3, ML210, FIN56, FINO2 and auranofin (for 24 h); erastin and sulfasalazine (for 48 h); BSO, iFSP1, viFSP1 and icFSP1 (for 72 h). Liproxstatin-1 (Lip-1, 0.5 µM) was used as a ferroptosis inhibitor. **(f)** Viability of various WT and *PRDX6* KO cancer cells treated with RSL3 (for 24 h) and erastin (for 48 h). Data is representative of 3 independent experiments. Data is mean ± s.d. of n = 3 (e) and mean of n = 3 (f).

### PRDX6 is involved in intracellular selenium metabolism

To systematically explore the contribution of PRDX6 to GPX4 expression, we sought to identify genetic co-dependencies using the depmap CRISPR screen database (**Fig 3a**). Among the top *PRDX6*-correlated genes, we found several genes crucial for the selenium metabolism pathway, such as LDL receptor-related protein 8 (*LRP8;* aka apolipoprotein E receptor 2*, APOER2),* selenophosphate synthetase 2 *(SEPHS2),* O-phosphoseryl-tRNA(Sec) selenium transferase *(SEPSECS)* and selenocysteine lyase (*SCLY*), alongside with genes involved in glutathione metabolism, such as glutathione synthetase (*GSS),* gamma-glutamylcysteine ligase, catalytic subuni*t (GCLC)* and gamma-glutamylcysteine ligase, modifier subunit *(GCLM*) and the cognate ferroptosis suppressor apoptosis inducing factor mitochondria-associated 2 (*AIFM2),* encoding FSP1 (**Fig. 3a**). Supporting this notion, a gene cluster analysis of coessentiality correlation network data^35^ also suggested the close relationship of *PRDX6* with other genes involved in selenium metabolism and selenoproteins including thioredoxin reductase 1 (*TXNRD1*) and *GPX4* (**Fig. 3b**) as previously reported^28^. In addition, a correlation of the expression level between PRDX6 and genes associated with the selenium metabolism pathway and other selenoproteins was also observed in the depmap proteomics dataset in cancer cell lines (**Fig. S3**). These correlations thus strongly suggest that PRDX6 may regulate GPX4 expression through selenium handling.

**Figure 3.**
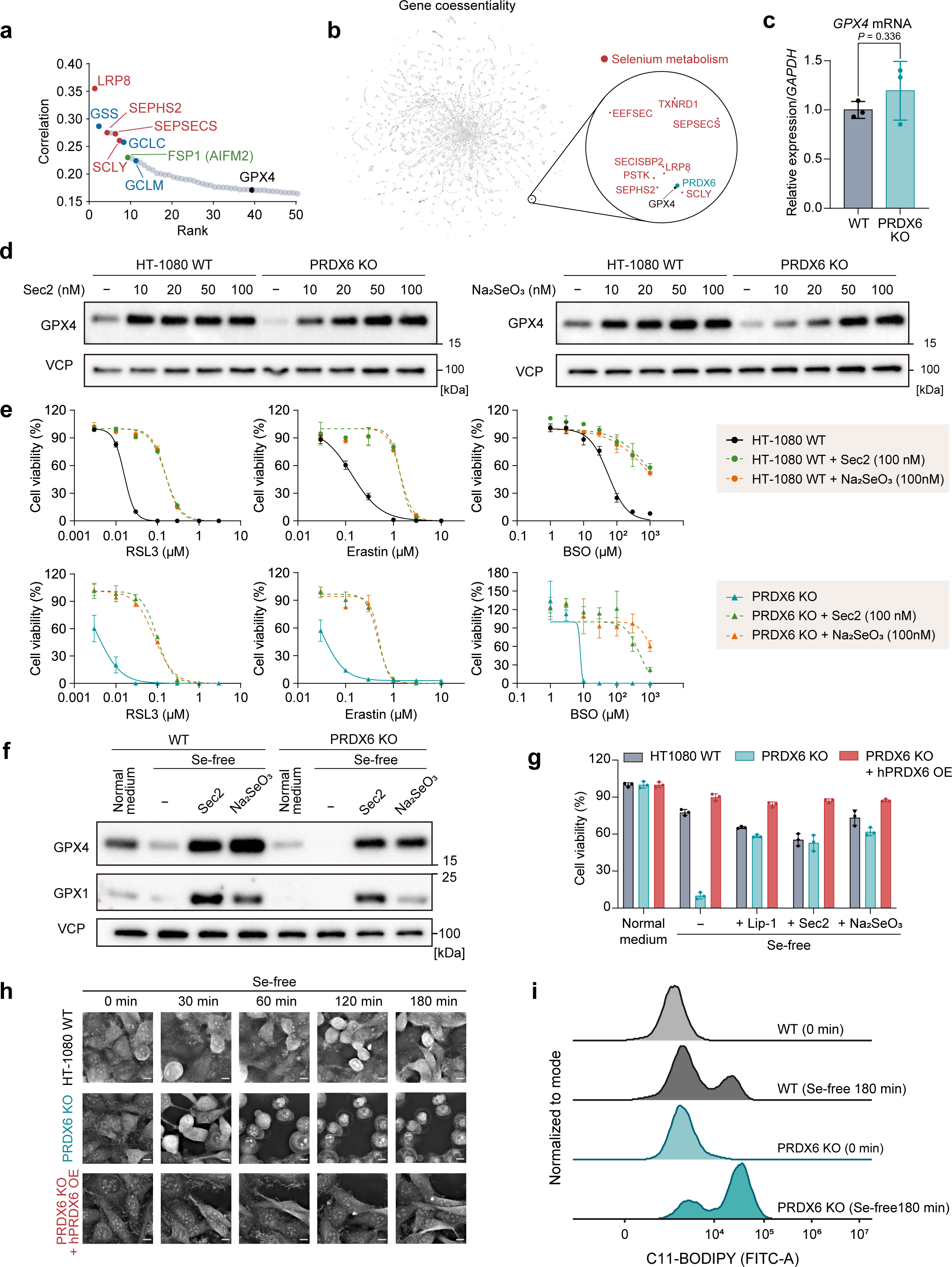
| PRDX6 is involved in selenoprotein synthesis and selenium metabolism **(a)** Top co-dependent genes with *PRDX6* in depmap CRISPR screen database (https://depmap.org; version v23Q4). Genes related to selenium metabolism (red), glutathione synthesis (blue) and ferroptosis suppressors (green) are highlighted. **(b)** Clustering genes from co-essentiality analysis (http://coessentiality.net). Selenium metabolism-related cluster is highlighted. **(c)** *GPX4* mRNA level in WT and *PRDX6* knockout (KO) HT-1080 cells. **(d)** Immunoblot analysis of GPX4 in WT and *PRDX6* KO HT-1080 cells treated with indicated concentrations of L-selenocystine (Sec2) or sodium selenite (Na_2_SeO_3_) for 48 h. **(e)** Viability of WT and *PRDX6* KO HT-1080 cells co-treated with ferroptosis inducers (RSL3, erastin and BSO) and different selenium sources (Sec2 or Na_2_SeO_3_). (f) Immunoblot analysis of GPX4 in WT and *PRDX6* KO HT-1080 cells after replacing normal culture medium with Se-free medium supplemented with Sec2 (100 nM) or Na_2_SeO_3_ (100 nM) for 48 h. (g) Viability of WT, *PRDX6* KO and *PRDX6* KO HT-1080 cells overexpressing hPRDX6 (OE) after replacing normal culture media with Se-free medium supplemented with Lip-1 (0.5 µM), Sec2 (100 nM) or Na_2_SeO_3,_ (100 nM). (h) Live cell imaging after replacing normal culture medium with Se-free medium. Scale, 10 µm. **(i)** C-11 BODIPY 581/591 lipid peroxidation assay of WT and *PRDX6* KO HT-1080 cells before (0 min) and after 180 min-incubation in Se-free media. Data represent the mean ± s.d. of n = 3 (c); the mean ± s.d. of 3 wells from one out of three independent experiments (e, g). t-test in (c).

Next, we found that *PRDX6* KO does not alter *GPX4* mRNA levels (**Fig. 3c**), suggesting that the impact of PRDX6 deletion on decreased GPX4 expression occurs at the translational level rather than at the transcriptional level. Since co-translational Sec incorporation at the UGA opal codon is the rate-limiting step in GPX4 translation^28,36^, we next examined the effect of selenium supplementation on the decreased level of GPX4 by *PRDX6* KO. When L-selenocystine (the dimeric and oxidized form of Sec; Sec2) or sodium selenite (Na_2_SeO_3_) were supplemented into the culture medium as selenium sources, the expression of GPX4 was increased in a dose-dependent manner starting at a 10 nM dose of each selenium source in *PRDX6* KO cells to the same extent as in WT cells (**Fig. 3d**). Both selenium sources also rescued the increased susceptibility to ferroptosis observed in *PRDX6* KO cells (**Fig. 3e**). These findings indicate that selenium supplementation can counteract the decrease in GPX4 expression and the resulting increase in ferroptosis vulnerability due to loss of *PRDX6*.

The effects of selenium deficiency were examined to further investigate the emerging relationship between selenium, PRDX6 and GPX4. When cells were cultured with selenium-deficient media for 48 h, GPX4 expression was decreased in WT cells, with almost no GPX4 expression being detectable in PRDX6 KO cells (**Fig. 3f**). A similar tendency was found for glutathione peroxidase 1 (GPX1), another selenoprotein family member known to swiftly react to changing selenium availability^37^. The reduction in GPX4 and GPX1 expression due to selenium deficiency was entirely restored by the supplementation of cells with L-selenocystine or sodium selenite (**Fig. 3f**). While WT HT-1080 cells remained viable under selenium-deficient conditions, *PRDX6* KO cells exhibited strongly decreased viability, which was entirely restored by Lip-1 and selenium supplementation as well as reconstitution of PRDX6 expression, demonstrating that selenium deprivation triggered ferroptosis in *PRDX6* KO cells (**Fig. 3g**). This selenium deprivation-induced cell death in *PRDX6* KO cells was began to be induced within several hours after replacing the standard medium with selenium-deficient medium (**Fig. 3h**). Consistently, staining of cells with the lipid peroxidation dye C11-BODIPY 581/591 assay showed that selenium deprivation increased lipid peroxidation in *PRDX6* KO cells in a similar time frame (**Fig 3i**). These results indicate that PRDX6 is involved in selenium metabolism and regulation of selenoprotein expression, including GPX4. However, *PRDX6* KO did not significantly affect the amount of total cellular selenium or the uptake of L-selenocystine and sodium selenite supplemented into the media (**Fig. S4**). This evidence suggests that PRDX6 acts downstream of selenium uptake, for instance, by influencing the efficiency of selenium utilization in cells.

### Cysteine 47 of PRDX6 is involved in selenium handling

To investigate the molecular basis of selenium metabolism by PRDX6, we focused on its different active sites relevant for its peroxidase activity (C47)^24^, PLA_2_ activity (S32)^23^ and dimer formation (C91)^38^ (**Fig. 4a**). To this end, we site-directedly mutated the respective residues in PRDX6 and stably expressed these variants in *PRDX6* KO HT-1080 cells. While the expression of S32A and C91S allowed the restoration of GPX4 and GPX1 expression levels to WT levels, the C47S mutant failed to restore the expression of both GPX4 and GPX1 (**Fig. 4b**). Accordingly, the increased ferroptosis sensitivity of *PRDX6* deficient cells to RSL3 and erastin was restored by overexpression of hPRDX6 WT, S32A and C91S mutants, but not the C47S variant (**Fig. 4c** and **4d**).

**Figure 4.**
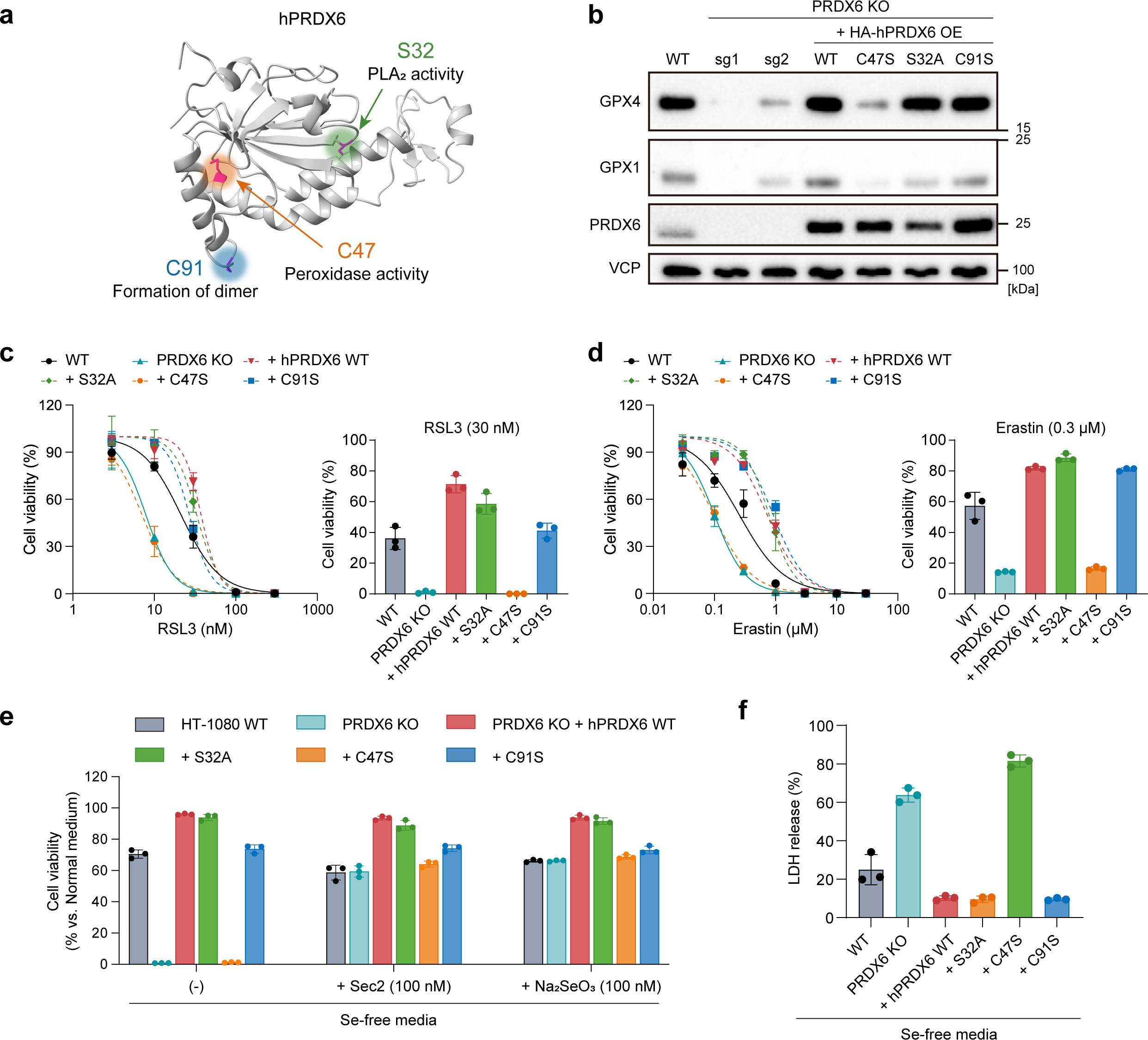
| PRDX6 C47 is critical for its role in selenium metabolism **(a)** A crystal structure of hPRDX6 (PDB:1PRX) highlighting the functional residues studied here. **(b)** Immunoblot analysis of WT HT-1080 cells, *PRDX6* KO (sg1 and sg2), *PRDX6* KO (sg2) overexpressing (OE) each of the different mutants of hPRDX6 (HA-tagged). **(c, d)** Viability of WT, *PRDX6* KO and *PRDX6* KO HT-1080 cells overexpressing each of the different hPRDX6 mutants treated with RSL3 and erastin for 24 h (left). Data for 30 nM of RSL3 and 0.3 µM of erastin is also shown as a bar graph (right). (e) Viability of cells 24 h after replacing the normal culture medium with Se-free medium supplemented with or without different selenium sources (100 nM of Sec2 or Na_2_SeO_3_). (f) LDH release level from cells cultured in a Se-free medium for 24 h. Data represents the mean ± s.d. of 3 wells from one out of three independent experiments (c-f).

Next, the different PRDX6 mutant lines were cultured in selenium-deficient media, revealing that only *PRDX6* KO cells and the cells expressing C47S variant succumbed to cell death, which could be rescued by supplementation of L-selenocystine or sodium selenite (**Fig. 4e** and **4f**). Of note, treatment with MJ33, which inhibits the PLA_2_ activity of PRDX6^39^, did not alter the susceptibility of HT1080 WT cells to ferroptosis, indicating that the PLA_2_ function is not involved in the regulatory action of PRDX6 in ferroptosis (**Fig. S4b**). These data shows that in the absence of abundant selenium, the peroxidative cysteine C47 in PRDX6 is essential for efficient intracellular selenium handling and proper expression of selenoproteins represented by GPX4, thereby protecting cells against ferroptosis.

### PRDX6 plays a role in efficient selenium mobilization

The general biosynthetic pathway for selenium metabolism for selenoproteins biosynthesis is summarized in **Fig. 5**^40^. Selenium uptake and intracellular metabolism can be broadly divided into an organic and inorganic pathway. In the organic pathway, the selenium transport protein selenoprotein P (SELENOP) is taken up by LRP8-mediated endocytosis and degraded in lysosomes whereupon Sec is released^28,41^. Sec is also supplied by the reduction of selenocystine (the oxidized dimeric form of Sec) that is taken up by xCT^42^. Subsequently, selenium is released from Sec in the form of selenide by SCLY^43^ or an SCLY-independent pathway^44^. In the inorganic pathway, selenite and its reduced form, selenide, are transported via different pathways and used as selenium sources^40^. Among the pathway, extracellular selenite is converted to its reduced form, selenide, in a step involving xCT (aka SLC7A11 and system x_c_^-^)^45,46^. xCT imports extracellular cystine, the oxidized dimeric form of cysteine (note, most cysteine is present in its oxidized form in medium) into cells. After conversion to its reduced form cysteine, it is used for GSH and protein biosynthesis, whereas a fraction is released into the extracellular space^47,48^. This xCT-mediated cystine-cysteine cycle provides extracellular thiol groups to reduce selenite to selenide. Eventually, intracellular selenide derived from both pathways is used as a substrate of SEPHS2 that catalyzes the production of monoselenophosphate, which is then used by SEPSECS to convert O-phosphoseryl-tRNA(Sec) to selenocysteinyl-tRNA(Sec), the latter being used by ribosomes for co-translational incorporation of Sec into selenoproteins^36^.

**Figure 5.**
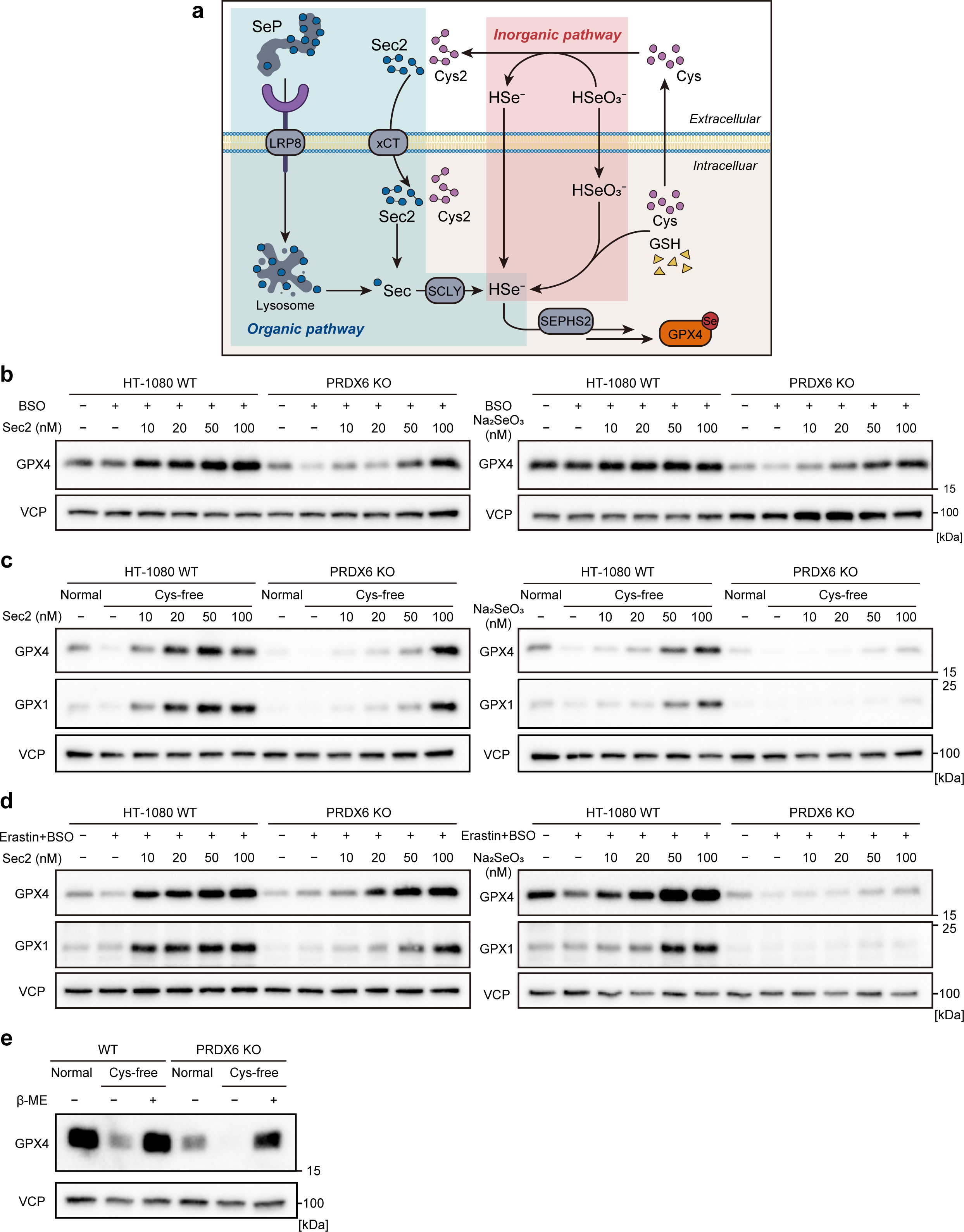
| PRDX6 is involved in cellular selenium mobilization **(a)** Scheme of the commonly known pathways for cellular selenium uptake and utilization for selenoprotein synthesis. Cys, cysteine; Cys2, cystine; Sec, selenocysteine; Sec2, selenocystine; SeP, selenoprotein P. The illustration was generated using BioRender.com. **(b)** Immunoblot analysis of WT and *PRDX6* KO HT-1080 cells treated with BSO (500 µM), Lip-1 (1 µM) and indicated concentrations of L-selenocystine (Sec2) or sodium selenite (Na_2_SeO_4_) for 72 h. **(c)** Immunoblot analysis of WT and *PRDX6* KO HT-1080 cells after replacing normal media with cyst(e)ine (Cys)-free media supplemented with 10% dialyzed FBS, Lip-1 (1 µM) and indicated concentrations of Sec2 or Na_2_SeO_3_ for 72 h. **(d)** Immunoblot analysis of WT and *PRDX6* KO HT-1080 cells treated with erastin (10 µM), BSO (500 µM), Lip-1 (1 µM) and indicated concentrations of Sec2 or Na_2_SeO_4_ for 48 h. (e) Immunoblot analysis of WT and *PRDX6* KO HT-1080 cells treated with Cys-free media with or without β-mercaptoethanol (β-ΜΕ, 50 µM) for 48 h.

The present studies suggested that PRDX6 is involved in selenium metabolism via its peroxidase activity, i.e., its reducing activity, at C47. Therefore, we assessed the involvement of PRDX6 on the expression of GPX4 under cell culture conditions with decreased reducing potential by depleting GSH and/or cysteine since the cysteine/xCT/GSH pathway generates the major cellular reducing capacity^49^. When cells were deprived from GSH by BSO, L-selenocystine and sodium selenite supplementation increased GPX4 expression both in WT and *PRDX6* KO cells, although *PRDX6* KO cells required a higher amount of selenium (50-100 nM) to afford an increased expression of GPX4 (**Fig. 5b**). When cells were cultured in cyst(e)ine free (Cys-free) media, which depletes both GSH and Cys^50^, expression of GPX4 and GPX1 was decreased in WT cells as previously reported^51^ and almost absent in *PRDX6* KO cells (**Fig. 5c**). The diminished expression of GPX4 and GPX1 in Cys-free media was rescued by supplementing either with L-selenocystine or sodium selenite in a dose-dependent manner in WT cells, whereas, interestingly, it was rescued in *PRDX6* KO cells by high concentration of L-selenocystine (100 nM), but hardly by sodium selenite supplementation even at concentrations up to 100 nM (**Fig. 5c**). The condition of the Cys-free media was pharmacologically reproduced with inhibitors by cotreatment of the cells with BSO and erastin (**Fig. 5d**). The results showed that the expression of GPX4 and GPX1 in *PRDX6* KO cells was rescued by the addition of L-selenocystine at higher concentrations than required for WT cells, whereas it was hardly increased by sodium selenite supplementation, similar to the result found in Cys-free media. These results suggest that in the absence of cellular reducing activity by depletion of Cys and GSH, KO of *PRDX6* renders selenite as an almost unusable source. Supporting this notion, treatment with a reducing agent, β-mercaptoethanol, restored decreased GPX4 expression in *PRDX6* KO cells cultured in Cys-free media (**Fig. 5e**). In summary, the results suggest that PRDX6 plays an important role in the efficient selenite utilization in concert with GSH and Cys.

## Discussion

In this manuscript, we revealed an unexpected role of PRDX6 in selenium metabolism. We showed that, beyond acting as a redox enzyme and phospholipase, PRDX6 facilitates cellular selenium mobilization, which is crucial for selenium incorporation into selenoproteins, including GPX4. Via this route, PRDX6 acts as a rheostat, modulating GPX4 expression level, thereby dictating the sensitivity of cells to undergo ferroptosis (**Fig. 6**).

**Figure 6.**
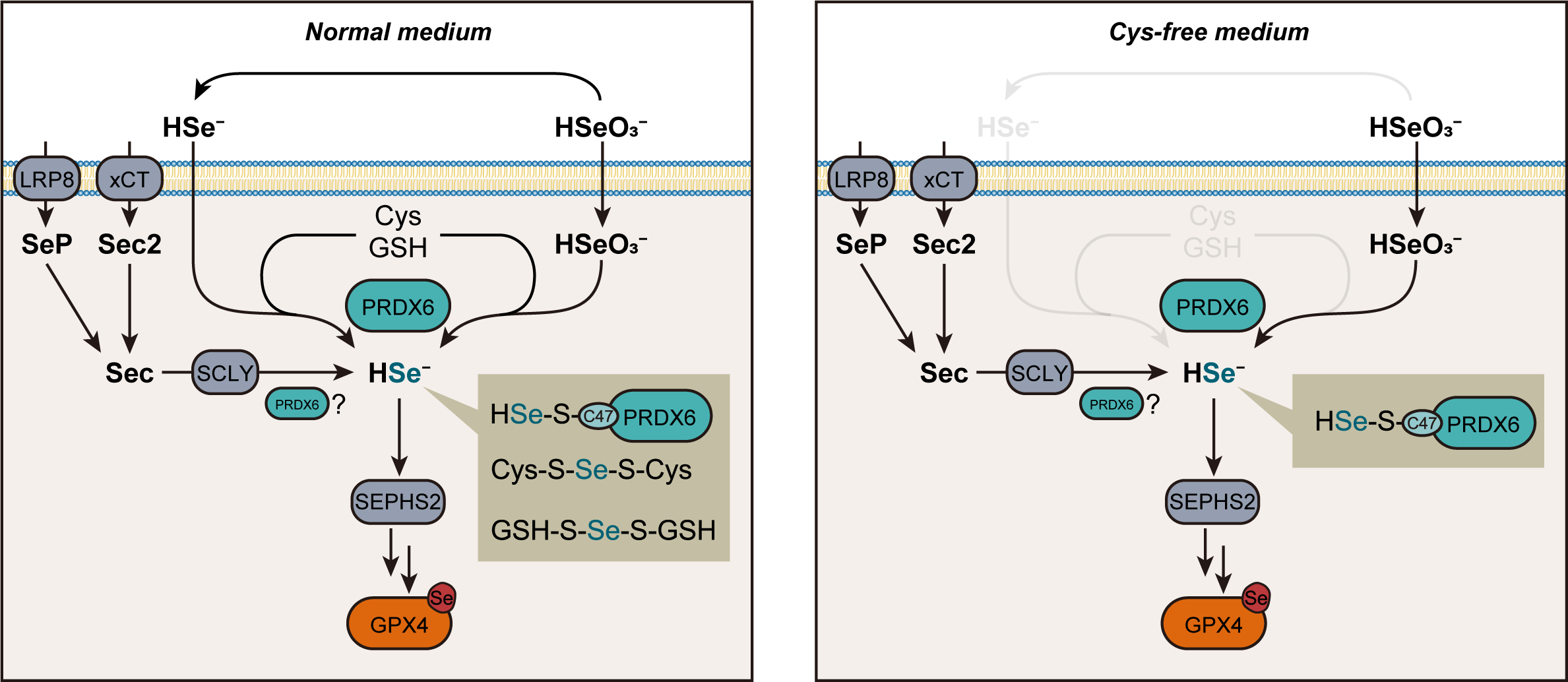
| PRDX6 as a crucial player in cellular selenium mobilization Proposed model for the role of RPDX6 in intracellular selenium handling. **(Left)** In the inorganic selenium pathway, selenite or its reduced form selenide is first taken up by cells. Cellular selenide is presumably bound to cysteine, GSH or PRDX6, which act as mobile selenium carriers. In the organic selenium pathway, selenide is generated from selenocysteine (Sec) derived from selenoprotein P (SeP) degradation or selenocystine (Sec2) taken up via xCT in a SCLY dependent- and independent-pathway. Selenide is eventually utilized as a selenium source for the biosynthesis of selenoproteins including GPX4. **(Right)** In cys(e)ine (Cys)-free medium conditions, in which Cys and GSH are depleted, extracellular selenite cannot be reduced to selenide due to the lack of the cystine/cysteine cycle, maintaining a fraction of selenite outside cells. Upon uptake, selenite is intracellularly reduced to selenide, which binds to PRDX6 for acting as a mobile selenium carrier. PRDX6-bound selenide then serves as the substrate for SEPHS2 for the conversion of O-phosphoseryl-tRNA(Sec) to selenocysteinyl-tRNA(Sec). While PRDX6-bound selenide can be utilized as a selenium source for the biosynthesis of selenocysteinyl-tRNA(Sec), in the *PRDX6* KO context, especially in Cys-free condition, selenite can hardly serve as a selenium source.

In particular, our finding that PRDX6-deficient cells can hardly utilize inorganic selenite as a selenium source in the absence of Cys and GSH highlights the orchestrating role of PRDX6 in cellular selenium metabolism. The catalytic residue C47 was found to be essential for the selenium handling function of PRDX6. Given the highly reactive nature of selenide (HSe^-^)^52^, a reduced form of selenite, it is postulated that its intracellular trafficking occurs by binding to acceptor molecules and/or proteins^43^, functioning as a “mobile selenium shuttle” within cells. Our findings suggest that PRDX6 is the carrier for selenium mobilization in cooperation with Cys and GSH, which have been reported to reduce selenite to selenide^46^. The reaction of GSH with selenite is known to form GSH-Se (*i.e.,* GSH-S-Se-S-GSH)^53^. In addition, a previous study has reported that C47 in PRDX6 is a molecular target for an electrophile^54^. Thus, mechanistically, it is plausible that PRDX6, using its reducing active site (-SH group), facilitates the process of the reduction of intracellular selenite to selenide and its efficient transfer to SEPHS2, enabling efficient selenium incorporation into selenoproteins (**Fig. 6**).

Our results further demonstrated that supplemented organic L-selenocysteine was still available in PRDX6-deficient cells, albeit requiring higher concentrations than in WT cells to increase sufficient GPX4 and GPX1 levels. This suggests that besides the role of PRDX6 in the inorganic selenium pathway, it also contributes to the efficient utilization of Sec in the organic pathway, comprising both SCLY-dependent and -independent pathways^44^. Thus, PRDX6 may also act as a selenide acceptor in these pathways. Previous studies have reported that cells employ distinct selenium uptake/utilization pathways based on their cell types^44^. Therefore, the impact of PRDX6 on selenium utilization and ferroptosis susceptibility may vary across cell types. Nevertheless, our findings consistently demonstrated the influence of *PRDX6* KO on GPX4 expression and ferroptosis sensitivity across a diverse range of cancer cells.

While both GPX4 and PRDX6 possess enzymatic activity in directly reducing PLOOH, our results suggest that they are unexpectedly not major contributors to overall PLOOH-reducing capacity in cells, as evidenced by considerable remaining PLOOH-reductive ability in GPX4 or PRDX6 deficient cells. Future research should, therefore, investigate the molecular identity responsible for cellular PLOOH reduction beyond GPX4 and PRDX6.

In aggregate, data presented by us and the accompanying manuscript by Chen et al. introduce PRDX6-dependent cues in selenium metabolism as critical checkpoints impacting ferroptosis surveillance^55^. On a more translational level, we showed that loss of *PRDX6* KO rendered cancer cells highly susceptible to ferroptosis, highlighting potential tractability in future anti-cancer drug design. Simultaneously, PRDX6 has been reported to be upregulated in neurodegenerative diseases^56,57^, which may instigate an investigation of this observation in the broader context of translational ferroptosis research.

### Limitations of the study

While our study proposes an important role for PRDX6 in ferroptosis regulation by impacting selenoproteins expression, we still need to understand its role *in vivo* as a selenium mobilizer. This is of particular relevance due to the greatly varying redox conditions and selenium sources between cell culture and a whole organism, thus requiring a careful validation of the present findings in *in vivo* studies.

## Methods

### Chemicals

Liproxstatin-1 (Lip-1, Selleckchem, S7699), erastin (Merck, 329600), (*1S,3R*)-RSL3 (RSL3, Cayman, 19288), iFSP1 (Cayman, 29483), ML210 (Cayman, 23282), FIN56 (Cayman, 25180), FINO2 (Cayman, 25096), L-buthionine sulfoximine (BSO, Sigma-Aldrich, B2515), auranofin (Sigma-Aldrich, A6733), sulfasalazine (Sigma-Aldrich, S0883), 4-hydroxytamoxifen (4-OH TAM, Sigma-Aldrich, H7904), L-selenocystine (Sigma-Aldrich, 545996), sodium selenite (Sigma-Aldrich, S5261), viFSP1 (Vistas-lab, STK626779), and MJ33 (Cayman, 90001844) were purchased. icFSP1 was synthesized by Intonation Research Laboratories.

### Cell culture

Murine immortalized 4-hydroxytamoxifen (TAM)-inducible *Gpx4* knockout (KO) fibroblasts (referred to as Pfa1) were reported previously^29^. Genomic *Gpx4* deletion can be achieved by TAM-inducible Cre recombinase using the CreERT2/LoxP system. HT-1080 (CCL-121), 786-O (CRL-1932), A375 (CRL-1619), A549 (CCL-185), HEK293T (CRL-3216) and NCI-H460 (HTB-177) cells were obtained from ATCC. LOX-IMVI cells were obtained from NCI/NIH. HEC151 cells (JCRB1122-A) were obtained from JCRB Cell Bank. Pfa1, HT-1080, 786-O, A375 and A549 cells were cultured in DMEM high glucose (4.5 g glucose/L) supplemented with 10% fetal bovine serum (FBS), 2 mM L-glutamine, and 1% penicillin/streptomycin. LOXIMVI, H460 and HEC151 cells were cultured in RPMI 1640 GlutaMax supplement medium (Gibco) with 10% FBS and 1% penicillin/streptomycin. For generating stably overexpressing cell lines, appropriate antibiotics (puromycin 1 µg/mL, blasticidin 10 µg/mL or G418 0.5-1 mg/mL) were used. *GPX4* KO HT-1080 and *Gpx4* KO Pfa1 cells were maintained in a medium containing Lip-1 (1 µM) to prevent ferroptosis. All cells were cultured at 37 °C with 5% CO_2_ and verified to be negative for mycoplasma.

### Preparation of PCOOH-*d*_9_

To obtain 1-palmitoyl-2-linoleoyl-sn-glycero-3-phosphatidylcholine-*d*_9_ (PC-*d*_9_) as a substrate of PCOOH-*d*_9_, 25 mg of 1-palmitoyl-2-linoleoyl-sn-glycero-3-phosphatidylethanolamine (16:0/18:2-PE, Avanti Polar Lipids, 850756) dissolved in methanol containing NaOH (1 N) was incubated with 250 µL of iodomethane-*d*_3_ (Isotec, 176036) for 6 h at 35 °C. After incubation, PC-*d*_9_ was isolated using preparative thin-layer chromatography and purified by solid phase extraction (Silica, Waters, WAT023595). To obtain PCOOH-*d*_9_, PC-*d*_9_ was subjected to photo-oxidation according to a previous study method with slight modifications^58,59^. Briefly, PC-*d*_9_ was dissolved in methanol containing 40 µM rose bengal, and then the sample was placed under the LED light and oxidized for 48 h at 40 °C. After the oxidation step, the sample was subsequently passed through a Sep-Pak NH_2_ cartridge (Waters, WAT023610) to remove rose bengal followed by evaporation and dissolving in methanol. Finally, PCOOH-*d*_9_ was isolated from crude oxidized PC-*d*_9_ using semipreparative LC.

### Measurement of PCOOH-*d*_9_ using LC-MS/MS

Cells were harvested and lysed by incubation with 0.1% Triton X-100 solution on ice for 30 min. Following centrifugation at 20,000 x g at 4 °C for 40 min, the supernatant was collected and used for PCOOH reduction assay. For evaluation of PCOOH-*d*_9_ reduction, 2 µL of PCOOH-*d*_9_ (10 µM in ethanol) was added to 198 µL of cell lysate and then incubated for 1 h at 37 °C. Then, PCOOH-*d*_9_ was extracted from 180 µL of the incubated sample by the Folch method according to a previously published method^60,61^ and dissolved in 200 µL of methanol. These methanol samples were subjected to an LC-MS/MS system. Data was analyzed using Analyst v1.7.2 software (Sciex). The detailed analytical conditions are described in Supplementary Information.

### Measurement of PCOOH in the liver of mice using LC-MS/MS

Frozen-stored tissues collected from *Alb-creERT2; Gpx4^fl/fl^* (hepatocyte-specific *Gpx4* KO) and *Cre(-); Gpx4^fl/fl^* (control) mice fed a low vitamin E diet (containing <7 mg/kg vitamin E, E15314-247, ssniff Spezialdiäten) for seven days after tamoxifen injection (2 mg on two consecutive days, dissolved in Miglyol 812, Caelo) were used. These tissues were the samples collected from the same mice used in the previous study^13^ and stored at −80°C. All experiments were performed in compliance with the German Animal Welfare Law and have been approved by the institutional committee on animal experimentation and the government of Upper Bavaria (approved no. ROB-55.2-2532-Vet_02-18-13). Total lipids containing PCOOH were extracted from the mouse liver samples by the Folch method^61,62^. PCOOH was analyzed using LC-MS/MS as described previously^63–65^. Data was analyzed using Analyst v1.7.2 software (Sciex). The detailed methods are described in Supplementary Information.

### Cell viability assay

Cells were seeded on 96-well plates and cultured overnight (2,000 cells/well for ferroptosis inducer treatment except for BSO and 4-OH TAM and 500 cells/well for BSO and 4-OH TAM treatment). On the next day, the medium was changed to the medium containing the compounds described in the corresponding figure legends at the indicated concentrations. After incubation for 24 h (RSL3, ML210, FIN56, FINO2 and auranofin), 48 h (erastin and sulfasalazine) and 72 h (BSO, iFSP1, viFSP1, icFSP1 and 4-OH TAM), cell viability was assessed using 0.004% resazurin. As readout, the fluorescence was measured at Ex/Em = 540/590 nm using a SpectraMax M5 microplate reader or Spectra Max iD5 (Molecular devices) with SoftMax Pro v7 (Molecular devices) after 4 h of incubation in a normal culture medium containing resazurin.

### Deprivation and supplementation of selenium

For selenium deprivation, cells were incubated with selenium-free (Se-free) medium (DMEM high glucose medium containing 2.5 mg/mL bovine serum albumin, 5 µg/mL insulin, 5 µg/mL transferrin, 92 nM FeCl_3_, 2 mM L-glutamine and 1% penicillin-streptomycin)^66^. For the immunoblotting assay, cells were harvested after incubation with Se-free medium supplemented Lip-1 (0.5 µM; to prevent ferroptosis) for 48-72 h. Cell viability was measured 24 h after replacing with Se-free media. For supplementation with L-selenocystine, sodium selenite or Lip-1, these were added to the Se-free medium at the timing of medium replacement.

### Cyst(e)ine and GSH deprivation

For cyst(e)ine (Cys) deprivation, cells were incubated with cystine/methionine-free high glucose DMEM (Gibco, 21013024) supplemented with 10% dialyzed FBS (Thermo Fisher, A3382001), 100 µM methionine, 1 mM sodium pyruvate, 2 mM L-glutamine and 1% penicillin-streptomycin. For the immunoblotting assay, cells were harvested 48-72 h after incubation in the Cys-free medium supplemented Lip-1 (0.5 µM) or β-mercaptoethanol (β-ΜΕ, 50 µM) to prevent ferroptosis. For BSO and/or erastin treatment, cells were harvested after incubation with normal culture media containing BSO (500 µM) and/or erastin (10 µM) with Lip-1 (0.5 µM; to prevent ferroptosis). L-selenocystine and sodium selenite were added at the same time when the medium was replaced.

### LDH release assay

HT-1080 cells (2,000 cells per well) were seeded on 96-well plates and cultured overnight. On the next day, the medium was changed to a selenium deprivation medium. Released LDH was assessed using the Cytotoxicity Detection kit (Roche, 11644793001) to determine necrotic cell death. In brief, cell culture supernatant was collected as a medium sample; then cells were lysed using 100 µL of 0.1% Triton X-100 in PBS as a lysate sample. Medium and lysate samples were individually loaded with reagents onto microplates, and the absorbance was measured at 492 nm using a SpectraMax M5 microplate reader after 15 min incubation at room temperature. LDH release (%) was calculated using medium sample values divided by the sum of medium and lysate sample values.

### Immunoblotting

Cells were lysed in LCW lysis buffer (0.5% Triton X-100, 0.5% sodium deoxycholate salt, 150 mM NaCl, 20 mM Tris-HCl, 10 mM EDTA and 30 mM sodium pyrophosphate tetrabasic decahydrate, pH 7.5) containing protease and phosphatase inhibitor mixture (cOmplete and phoSTOP; Roche, 04693116001 and 4906837001) and centrifuged at 20,000g for 40 min at 4 °C. After addition of 6 x SDS sample buffer (375 mM Tris-HCl, pH 6.8, 9% SDS, 50% glycerol, 9% β-mercaptoethanol and 0.03% bromophenol blue) to the collected supernatant and heating at 55 °C for 3 min, the samples were resolved on 12% SDS-PAGE gels and electroblotted onto a PVDF membrane (Bio-Rad, 170-4156). The membrane was blocked with 5% skim milk (Carl Roth, T145.2) in TBS-T (20 mM Tris-HCl, 150 mM NaCl and 0.1% Tween-20) and then probed with the primary antibodies against GPX4 (1:1,000, ab125066, Abcam), GPX1 (1:1,000, ab22604, Abcam), PRDX6 (1:1,000, 95336S, Cell Signaling Technology), PRDX1 (1:1000, rabbit, kindly provided by Dr. Christopher Horst Lillig, Greifswald, and Carsten Berndt, Düsseldorf)^67^, PRDX2 (1:1000, rabbit, provided by Christopher Horst Lillig, Greifswald, and Carsten Berndt, Düsseldorf)^68^, PRDX3 (1:1,000, Abcam, ab129206), PRDX4 (1:1,000, Abcam, ab184167), PRDX5 (1:1,000, Abcam, ab180587), FSP1 (1:1000, Santa Cruz, sc-377120), HA (1:1,000, clone 3F10, rat IgG1, developed in-house) and valosin containing protein (VCP, 1:10,000, ab11433, Abcam). Images were analyzed with Image Lab 6.0 software (Bio-Rad).

### Lentiviral production and transduction

A lentiviral transduction system was used for generating KO and overexpressing cells. A third-generation lentiviral packaging system consisting of transfer plasmids of the interest, pMD2.G (Addgene, 12259) and psPAX2 (Addgene, 12260) were co-lipofected into HEK293T cells using PEI MAX (Polysciences, 24765). Supernatants containing viral particles were harvested at 2 days post-transfection and filtered through a 0.45 µm PVDF filter (Millipore, SLHV033RS) and stored at −80 °C until use. Cells were seeded on 12 well plates with lentivirus particles and protamine sulfate (10 µg/mL) to enhance the transduction efficiency. On the next day, the cell culture medium was replaced with fresh medium containing respective selection antibiotics and cultured until non-transduced cells were dead.

### Generation of knockout cells

*Gpx4* KO Pfa1 cells were established by single cloning of 4-OH TAM-treated Pfa1 cells in the previous study^32^. *GPX4* KO HT-1080 cells and *GPX4* KO HT-1080 cells overexpressing hGPX4 were made in the previous study^69^. *PRDX6* and *PRDX1-5* KO cells were established by lentiviral transduction. Single guide RNAs (sgRNA) were designed to target exons of the genes of interest with the VBC score (https://vbc-score.org/) and the sequences are listed in Key Resource Table. The sgRNAs were subcloned into *BsmB*I-digested lentiCRISPRv2-puro vector (Addgene, 98290). One day after the transduction of lentiviral particles containing the transfer plasmid of lentiCRISPRv2-puro inserted sgRNA sequences, the cell culture medium was replaced with media containing puromycin (1 µg/ml). After selection for 2-3 days, polyclonal KO cells were subjected to the experiments. For the establishment of *PRDX6* KO HT-1080 cells, single-cell clones were isolated by serial dilution and knockout clones were validated by immunoblotting.

### Cloning of plasmids for overexpression

A human *PRDX6* cDNA (NM_004905.3) was cloned from the cDNA of HT-1080 cells and followed by cloning in the p442-blast vector furnished with an N-terminal HA tag. To avoid being targeted by CRISPR-Cas9, sgRNA-resistant mutants were created by site-directed mutagenesis using polymerase KOD One (Sigma, KMM-201NV) followed by ligation using in-Fusion cloning enzymes (Takara Bio, 638948). hPRDX6 mutants (S32A, C47S and C91S) were generated by site-directed mutagenesis. Codon-optimized *Mus musculus* (mouse) *PRDX6* (NP_031479.1) and *Homo sapiens* (human) GCH1 (NP_000152.1) were synthesized by Twist Bioscience and cloned into the lentivirus plasmid pLV-Neo. Human *FSP1* (NM_001198696.2) and short form of human GPX4 (NM_001367832.1) cloned into 442-blast were prepared in the previous studies^34,69^.

### Live-cell imaging

HT-1080 cells (6 x 10^4^ cells/well) were seeded on μ-Dish 35 mm low (80136, i-bidi) and incubated overnight. On the next day, live-cell imaging was performed using 3D Cell Explorer and Eve software v1.8.2 (Nanolive). During imaging, the cells were maintained at 37 °C and 5% CO_2_ by using a temperature-controlled incubation chamber.

### Lipid peroxidation assay

HT-1080 cells (1 x 10^5^ cells per well) were seeded on 12-well plates one day before the experiments. On the next day, the cell culture medium was replaced with selenium deprivation media for 180 min and subsequently incubated with C11-BODIPY 581/591 (1.5 µM, Invitrogen, D3861) for 30 min in a 5% CO_2_ atmosphere at 37°C. The cells were washed with PBS, trypsinized and resuspended in 500 µL PBS. After passing through a 40 μm cell strainer, cells were analyzed on a flow cytometer (CytoFLEX, Beckman Coulter) equipped with a 488 nm laser for excitation. Data was collected using CytExpert v2.4 (Beckman Coulter) from the FITC detector (oxidized BODIPY) with a 525/40 nm bandpass filter. At least 10,000 events were analyzed per sample. Data was analyzed using FlowJo Software (v10, FlowJo LLC).

### Structural data

The structure of the PRDX6 (PDB: 1PRX) is displayed using ChimeraX (https://www.cgl.ucsf.edu/chimerax/)70 for highlighting key functional residues.

### Database analysis of proteomics and CRISPR KO screening data

Proteomics data and CRISPR co-dependency of *PRDX6* were minded from depmap (http://www.depmap.org; version v23Q4). Co-essentiality analysis was performed using Co-essentiality browser (http://coessentiality.net/; date obtained in 2024.01).

### qPCR analysis

Total RNA was extracted from cells with RNeasy Mini kit (Qiagen, 74104) and followed by genomic DNA digestion using RNase-free DNase (Qiagen, 79254). cDNA was synthesized by QuantiTect Reverse Transcription Kit (Qiagen, 5001473). Quantitative RT-PCR (qPCR) was carried out by PowerUp SYBR Green Master Mix (Thermo Fisher, A25778) using qTOWER3 G (Analytikjena). All samples were performed with technical triplicates with the following PCR condition: [1] 50 °C for 2 min; [2] 95 °C for 2 min; [3] 95 °C for 15 sec; [4] 59.5 °C for 15 sec; [5] 72°C for 1 min; [6] 95°C 1 sec - cycle from 3 to 5 was repeated 40 times. Sequences of the primers were the following: 5’-TGCTCTGTGGGGCTCTG −3’ and 5’-ATGTCCTTGGCGGAAAACTC −3’ for short form GPX4; and 5’-GGTGTGAACCATGAGAAGTATGA −3’ and: 5’-GAGTCCTTCCACGATACCAAAG-3’ for GAPDH. The relative GPX4 expression was normalized to GAPDH by the ΔΔCT method.

### Measurement of cellular selenium level

To measure total cellular selenium levels, WT and *PRDX6* KO HT-1080 cells (2 x 10^5^ cells per well) were seeded on 6-well plates with standard culture medium. The following day, the medium was replaced with either standard culture medium containing Lip-1 (1 μM) or cyst(e)ine-free medium containing Lip-1 (1 μM) with or without supplementation of L-selenocystine (100 nM) or sodium selenite (100 nM). After 24 h of incubation, the cells were washed with PBS and harvested in 200 µL of PBS. The cells were separated into two aliquots and collected by centrifugation (1000 x g, 3 min, 4°C). One aliquot was solubilized with 1% SDS to determine protein content and the other aliquot was mixed with 1 mL 30% HNO_3_ and subjected to ICP/MS 8900 (Agilent). Data was normalized by the corresponding protein concentration.

### Statistical analysis

Statistical information for individual experiments can be found in the corresponding figure legends. Graphs were created using GraphPad Prism v10 (GraphPad Software).

## Acknowledgements

We thank Dr. Carsten Berndt (Heinrich Heine University) and Dr. Christopher Horst Lillig (University Medicine Greifswald) for kindly providing antibodies against PRDX1 and PRDX2; Dr. Sho Kobayashi (Yamagata University) and Dr. Ryuta Tobe (Tohoku University) for providing expert views. This work was supported by Deutsche Forschungsgemeinschaft (DFG) (CO 291/7-1, the Priority Program SPP 2306 [CO 291/9-1, #461385412; CO 291/10-1, #461507177] and the CRC TRR 353 (CO 291/11-1; #471011418), the German Federal Ministry of Education and Research (BMBF) FERROPATH (01EJ2205B) and the European Research Council (ERC) under the European Union’s Horizon 2020 research and innovation programme (grant agreement No. GA 884754) to M.C. and JSPS KAKENHI (20KK0363 and 18K08198 to E.M.; 22KK0253 to J.I. and 22H02278 to J.I. and K.N.).

## Author Contributions

J.I., T.N., E.M. and M.C. conceived experiments. J.I., T.N., T.T., M.S., W.Z., J.Z., N.Y. and K.O. performed experiments. J.I., T.N., E.M., A.W. and M.C. wrote the manuscript. T.D. performed data analysis. T.T., S.D., Y.S. and K.N. provided expertized feedback. J.I., E.M., K.N. and M.C. secured funding.

## Supplementary figure legends

**Figure S1 | Synthesis of PCOOH-*d*_9_.**

Scheme for the synthesis of deuterium-labelled phosphatidylcholine hydroperoxide (PCOOH-*d*_9_).

**Figure S2 | PCOOH-reducing capacity assay using Pfa1 cells and immunoblotting analysis of *GPX4* KO cells**

**(a)** Immunoblot analysis of HT-1080 WT and *GPX4* KO cells.

**(b)** PCOOH reducing capacity was measured using cell lysate collected from WT, *Gpx4* KO and *Prdx6* KO Pfa1 cells. *Gpx4* KO Pfa1 cells were established by single cloning of 4-hydroxytamoxifen-treated Pfa1 cells. The chromatogram (left) and relative value of the remaining PCOOH-*d*_9_ (right) are shown. Data is mean ± s.d. of n = 3.

**(c)** Immunoblot analysis of HT-1080 WT, *PRDX6* KO and *PRDX6* KO cells overexpressing hPRDX6.

**Figure S3 | Correlation of PRDX6 with selenoproteins and seleno-metabolism-related proteins.**

Correlation of PRDX6 with TXNRD1, SELENOH, SELENOW, SCLY and SEPHS2 in proteomics data sets (depmap; version v23Q4; http://www.depmap.org) is shown.

**Figure S4 | *PRDX6* KO does not affect cellular selenium uptake.**

**(a)** The total cellular selenium level of WT and *PRDX6* KO HT-1080 cells supplemented with L-selenocystine (Sec2, 100 nM) or sodium selenide (Na_2_SeO_3_, 100 nM) for 24 h. Cellular selenium level was measured by ICP/MS. Mean ± s.d. of n = 3 in each group. t-test.

**(b)** The effect of MJ33 (an inhibitor of the PLA2 activity of PRDX6) on RSL3-induced ferroptosis in HT-1080 WT cells. The cells were co-treated with RSL3 and MJ33 for 24 h. Data represents the mean ± s.d. of n = 3.

