## Supplementary figures and images for "PRDX6 dictates ferroptosis sensitivity by directing cellular selenium mobilization"

### Supplemental Figures

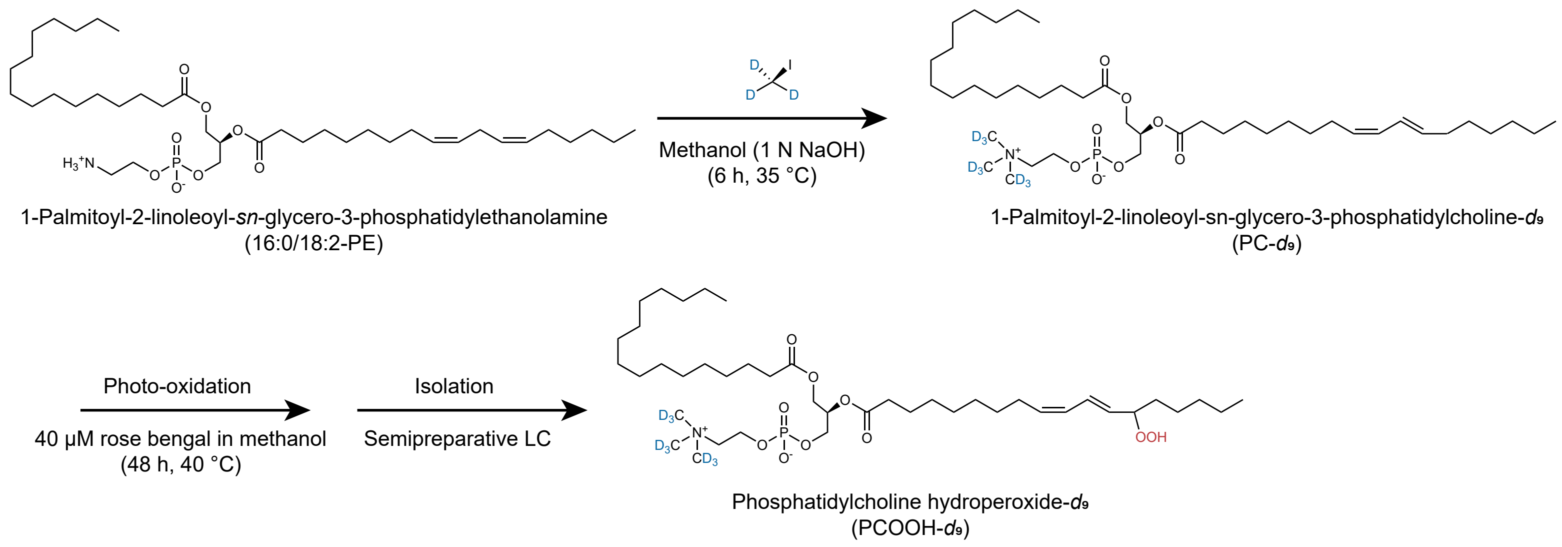

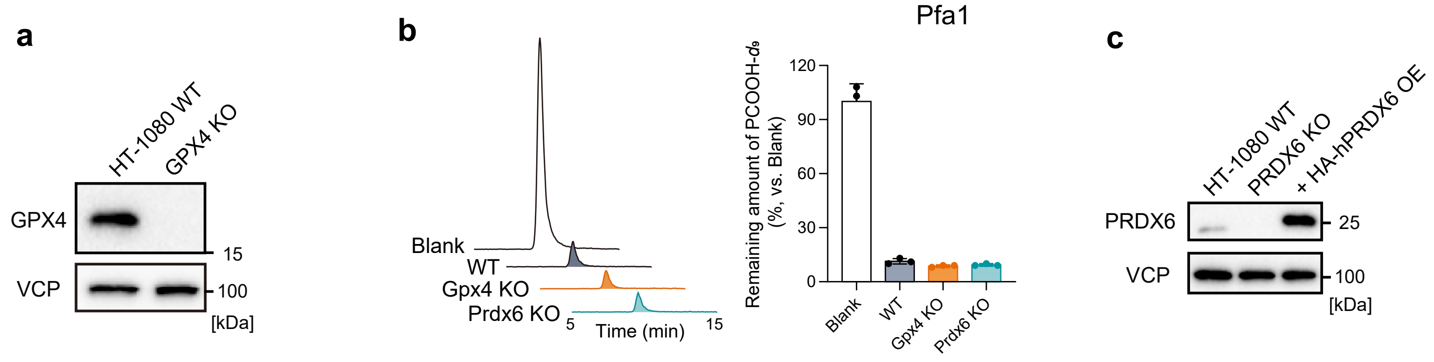

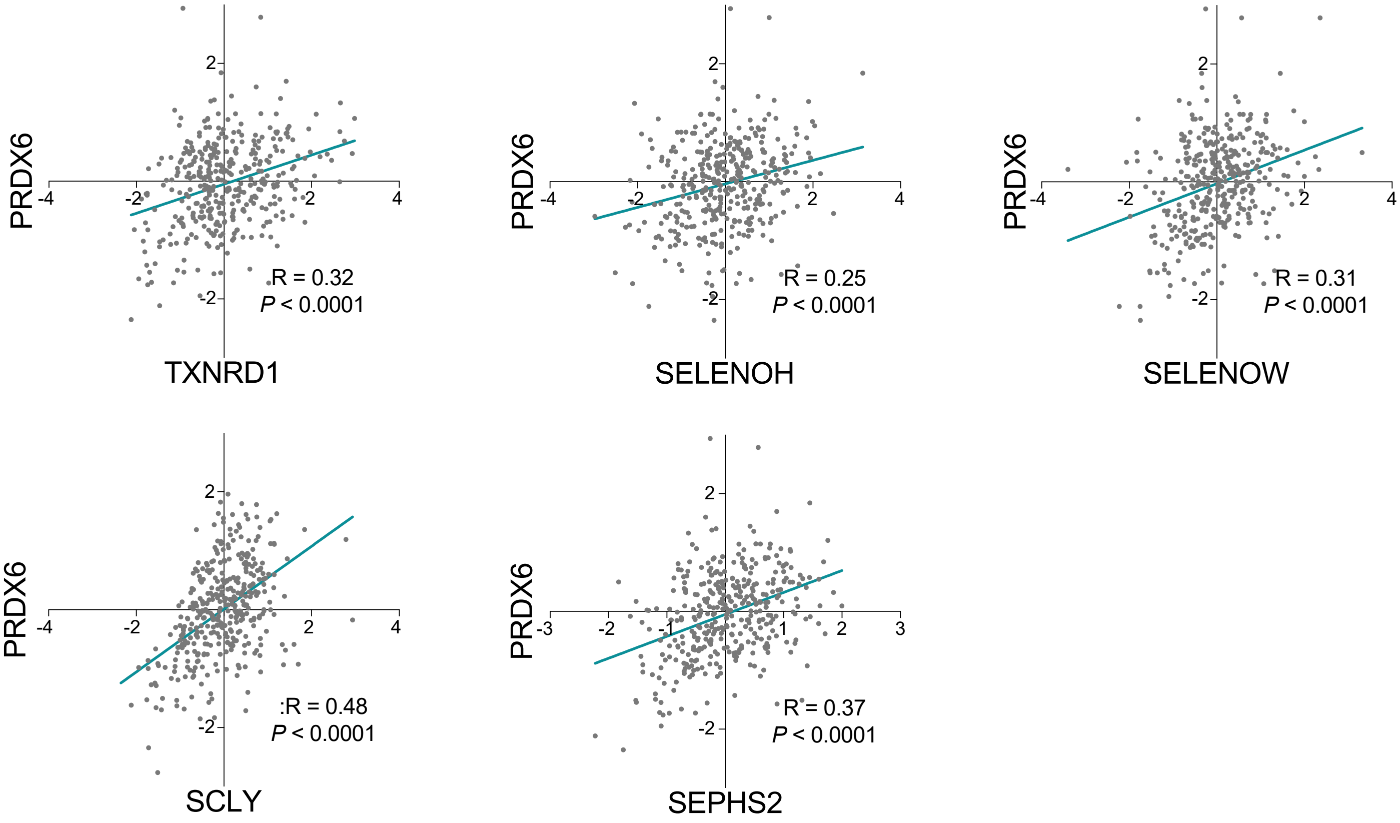

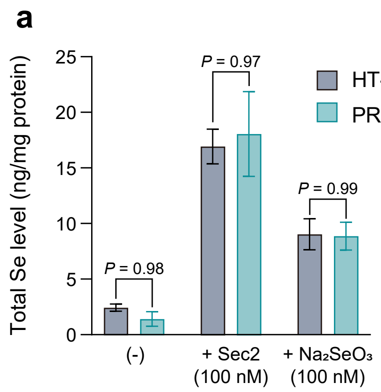

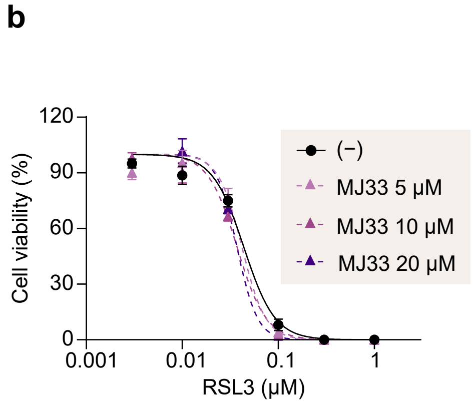
